# Relationship between systolic blood pressure and decision-making during emotional processing

**DOI:** 10.1101/2021.10.22.465429

**Authors:** Sophie Betka, David Watson, Sarah Garfinkel, Gaby Pfeifer, Henrique Sequeira, Theodora Duka, Hugo Critchley

## Abstract

**Objective:** Emotional states are expressed in body and mind through subjective experience of physiological changes. In previous work, subliminal priming of anger prior to lexical decisions increased systolic blood pressure (SBP). This increase predicted the slowing of response times (RT), suggesting that baroreflex-related autonomic changes and their interoceptive (feedback) representations, influence cognition. Alexithymia is a subclinical affective dysfunction characterized by difficulty in identifying emotions. Atypical autonomic and interoceptive profiles are observed in alexithymia. Therefore, we sought to identify mechanisms through which SBP fluctuations during emotional processing might influence decision-making, including whether alexithymia contributes to this relationship.

**Methods:** Thirty-two male participants performed an affect priming paradigm and completed the Toronto Alexithymia Scale. Emotional faces were briefly presented (20ms) prior a short-term memory task. RT, accuracy and SBP were recorded on a trial-by-trial basis. Generalized mixed-effects linear models were used to evaluate the impact of emotion, physiological changes, alexithymia score, and their interactions, on performances.

**Results:** A main effect of emotion was observed on accuracy. Participants were more accurate on trials with anger primes, compared to neutral priming. Greater accuracy was related to increased SBP. An interaction between SBP and emotion was observed on RT: Increased SBP was associated with RT prolongation in the anger priming condition, yet this relationship was absent under the sadness priming. Alexithymia did not significantly moderate the above relationships.

**Conclusions:** Our data suggest that peripheral autonomic responses during affective challenges guide cognitive processes. We discuss our findings in the theoretical framework proposed by Lacey and Lacey (1970).

## 1. Introduction

Emotional feeling states originate in part through subjective experience and cerebral representation of peripheral physiological reactions to affective stimuli (James, 1884; Lange, 1912). Bodily changes also inform and influence perceptual experience, allocation of attentional resources, emotional processing and decision-making (Bechara, Tranel, Damasio, & Damasio, 1996; Garfinkel et al., 2016; Gray, Minati, Paoletti, & Critchley, 2010; Makovac et al., 2017; Makovac et al., 2015; H. D. Park, Correia, Ducorps, & Tallon-Baudry, 2014; Łukowska, Sznajder, & Wierzchoń, 2018). One experimental means of testing how autonomic changes influence cognition is the use of an affective priming paradigm. Here, the emotional valence of a briefly presented stimulus (i.e. the prime) affects the processing of subsequent stimuli. Typically, if the prime and the target are of the same valence, a facilitatory effect (e.g. reduced reaction times) is observed. If the valence is different, an inhibitory effect is observed (e.g. increased reaction times) relative to a non-valenced control condition. In some forms of this paradigm, the prime is followed by a stimulus that prompts performance of a task, e.g. making a lexical decision (Hull, Slone, Meteyer, & Matthews, 2002). For example, the subliminal presentation of the word “ANGER” (vs “RELAX”) as a prime, just prior to rapid judgments of letter-strings, increases systolic blood pressure in healthy individuals. Here, the magnitude of this increase predicts RT prolongation on the task (Garfinkel et al., 2016). Increased systolic blood pressure is also observed in priming studies using emotional faces (Gendolla & Silvestrini, 2011; Lasauskaite, Gendolla, & Silvestrini, 2013; Silvestrini & Gendolla, 2011b, 2011c) (for review see van der Ploeg, Brosschot, Versluis, & Verkuil, 2017). Affect primes are proposed to activate affective mental representations influencing physiological reactivity (Gendolla, 2012), informing behavioral responses *via* afferent (interoceptive) feedback. However, in some forms of emotional dysfunction including alexithymia, the representation, and integration of interoceptive bodily responses appears to be impaired.

Alexithymia is a personality construct classically defined by difficulties in describing and identifying ones’ own emotional feelings (Apfel & Sifneos, 1979; Taylor, Ryan, & Bagby, 1985). Alexithymic subjects show abnormal affective regulation characterized by reduced emotional faces recognition, delayed automatic rapid facial reactions, reduced empathy, reduced emotional awareness, abnormal emotional remapping and higher body representation malleability (Georgiou, Mai, & Pollatos, 2016; Grynberg et al., 2012; Grynberg & Pollatos, 2015; Lane, Hsu, Locke, Ritenbaugh, & Stonnington, 2015; Moriguchi et al., 2007; Scarpazza, Ladavas, & Cattaneo, 2017; Scarpazza, Ladavas, & di Pellegrino, 2015).

Recent work suggests that interoceptive failure (e.g. reduced sensitivity to bodily sensations) is core to emotional impairments observed in alexithymia (Betka et al., 2017; Bird et al., 2010; Brewer, Cook, & Bird, 2016; Hogeveen, Bird, Chau, Krueger, & Grafman, 2016; J. Murphy, Catmur, & Bird, 2017; Shah, Hall, Catmur, & Bird, 2016). Indeed, visceral arousal informs the complex experience of subjective feelings (Critchley & Harrison, 2013; Shah, Catmur, & Bird, 2017; Tsakiris & Critchley, 2016). For example, one channel communicating cardiovascular arousal originates in afferent signals from arterial baroreceptors localized in the aorta and carotids. The arterial baroreflex is a homeostatic mechanism that minimizes fluctuation in blood pressure through coupling afferent baroreceptor signals to efferent control of heart rate, cardiac output and peripheral resistance (Brading, 1999; Eckberg & Sleight, 1992). Observations in hypertensive patients can inform our understanding of how interoceptive abnormalities may underlie the expression of alexithymia. In hypertension, abnormalities of the baroreflex mechanism are observed, including reduced baroreflex sensitivity (Mussalo et al., 2002). Hypertensive patients are also reported to show impaired interoceptive abilities during a heartbeat detection task (Yoris et al., 2017).This deficit is observed independently of heart rate or heart rate variability (HRV; an index of parasympathetic cardiac control) and exteroceptive abilities remain preserved (Yoris et al., 2017).This disruption of interoceptive processing in hypertensive patients is likely to contribute to the association between raised resting systolic blood pressure and impaired emotional processing (McCubbin et al., 2011; Pury, McCubbin, Helfer, Galloway, & McMullen, 2004).These findings suggest important significance of blood pressure on aspects of interoception.

Conversely, peripheral autonomic abnormalities are documented in alexithymic subjects who are at increased risk of premature death and cardiovascular disease (Helmers & Mente, 1999; Kauhanen, Kaplan, Cohen, Julkunen, & Salonen, 1996; Kauhanen, Kaplan, Cohen, Salonen, & Salonen, 1994; Tolmunen, Lehto, Heliste, Kurl, & Kauhanen, 2010). Dampened autonomic reactivity to emotional challenges or stress supports a hypoarousal model of alexithymia (Cecchetto, Korb, Rumiati, & Aiello, 2018; Constantinou, Panayiotou, & Theodorou, 2014; Neumann, Sollers, Thayer, & Waldstein, 2004; Peasley-Miklus, Panayiotou, & Vrana, 2016; Pollatos et al., 2011). However, there is yet no conclusive evidence for an association between alexithymia and abnormal baroreflex sensitivity (Virtanen et al., 2003). Alexithymic individuals report higher self-reported anxiety and show greater systolic blood pressure reactivity during the stress of blood donation (Byrne & Ditto, 2005). During anger recall, alexithymic individuals have attenuated cardiac responses (Neumann et al., 2004) but then show hampered recovery (of diastolic blood pressure recovery and cardiac preejection period), compared to non-alexithymic individuals (Neumann et al., 2004). Nevertheless, alexithymia is more prevalent among hypertensive patients than in the general population (Gage & Egan, 1984; Jula, Salminen, & Saarijarvi, 1999; Todarello, Taylor, Parker, & Fanelli, 1995).

A few studies have tested affective priming effects in alexithymia, but with variable methodologies. Priming by verbal stimuli is reportedly enhanced in alexithymia (Suslow, 1998), and greatest for negative primes when emotionally congruent with targets (Suslow, Junghanns, Donges, & Arolt, 2001). In a third study using different types of prime/target associations (e.g. verbal-verbal; facial-verbal; verbal facial etc.), alexithymia also reportedly moderates the impact of angry face primes on the evaluative judgment of an affective target, the effect apparent across verbal, visual and cross-modal trials (Vermeulen, Luminet, & Corneille, 2006). When primes and targets are specifically related to illness (e.g. verbal prime: “dizziness”), alexithymia was associated with faster reaction times for negative and positive verbal primes to illness-related targets (Brandt, Pintzinger, & Tran, 2015). These studies do not report implicit physiological responses.

### 1.1. Aims and hypotheses

Therefore, we sought to identify the relationship between physiological changes evoked by emotional primes (indexed by systolic blood pressure) and cognition (indexed by accuracy and speed of decision-making). A second aim was to explore whether alexithymia contributed to this relationship.

We expected that greater accuracy and shorter reaction times on a short-term / working memory task will be predicted by increased systolic blood pressure in the emotional priming conditions (anger and sadness) compared to the neutral priming condition.

Also, we hypothesized that alexithymia may partly mediate the relationship between physiological changes and decision-making through its association with increased systolic blood pressure at rest.

Given the well documented co-occurrence of depression, anxiety and alcohol use disorders in alexithymia (Finn, Martin, & Pihl, 1987; Hendryx, Haviland, & Shaw, 1991), we additionally assessed these variables in order to control for potential cofounding effects.

## 2. Methods

### 2.1. Participants

Thirty-two male volunteers (mean age 25.1 yrs; range 18–36yrs) took part in the experiment, based on a sample size calculation using statistical information from a similar study (Garfinkel et al., 2016). The present study contributed to the baseline session of a larger project looking at the impact of intranasal oxytocin on emotional regulation processes. For this reason, only male participants were recruited, via poster advertisements placed around the University of Sussex and Brighton and Sussex Medical School. All participants were healthy, with no history of psychiatric or neurological diseases. The average number of years of education was M= 16.9 (SD = 2.6). All participants gave their written informed consent and were compensated £ 7 per hour for their time. The study was reviewed and approved by the BSMS Research Governance and Ethics committee.

### 2.2. Material

### 2.3. Stimuli

A modified version of a Sternberg Task (e.g. item-recognition paradigm involving short-term / working memory) was designed(Sternberg, 1966). Ninety-six strings of seven letters were selected (e.g. KOPLTFV, IZTNLDS). Each presentation of a letter-string was followed by visual mask then the presentation of a target letter. The target letter was present in half of the letter-string trials. During the experiment, the letter-string was preceded by a visual prime: a very short presentation of an image displaying an emotional facial expression (EFE) of sadness, anger or neutrality. For each of these affective conditions, there were 32 trials (32 letter-strings preceded by an EFE of sadness, 32 letter-strings preceded by an EFE of anger, 32 letter-strings preceded by an EFE of neutrality). The EFE, coloured and from front perspective, were chosen from the Karolinska Directed Emotional Faces battery (Goeleven, De Raedt, Leyman, & Verschuere, 2008, for list of stimuli codes see the supplementary section). Trials were presented in blocks of four consecutive trials of the same EFE, balanced and randomised for male/female gender. This cognitive task was run using Matlab.

### 2.4. Physiological recording

Blood pressure was recorded using non-invasive, beat-to-beat monitoring via photoplethysmographic technology (Finometer PRO, Finapres medical systems, Amsterdam, The Netherlands). An inflatable finger cuff and infrared plethysmograph were fitted to the middle finger of the participant’s left hand, allowing measurements of beat-to-beat systolic blood pressure. The heart level electrode was attached to participant’s clothing in the mid-clavicular line at the level of the heart. Physiological data was recorded using Spike software 2 6.08.

### 2.5. Questionnaires

#### 2.5.1. Toronto Alexithymia Scale-20 items (TAS-20)

The TAS-20 (Bagby, Parker, & Taylor, 1994) consists of 20 items rated on a five-point Likert scale (from 1 “strongly disagree” to 5 “strongly agree”). Exploratory factor analysis and confirmatory factor analyses demonstrated that TAS-20 had a good internal consistency (Cronbach’s α=0.81), a good test-retest reliability (0.77, p <0.01) and a three-factor structure, in both clinical and non-clinical populations (Haviland, Shaw, MacMurray, & Cummings, 1988; Taylor et al., 1988). The TAS-20 is composed of three factors. The first factor measures difficulties in identifying feelings (DIF), the second factor measures difficulties in describing feelings (DDF) and the third factor measures the way the participant uses externally oriented thoughts (EOT). The total alexithymia score is the sum of responses across all 20 items. We only considered the total score in our analyses. High alexithymic are characterised by a score > or = 61; non alexithymic are characterised by a score < or = 51; between 50 and 60, subjects are characterised as intermediate.

#### 2.5.2. Trait Anxiety (STAI)

Trait anxiety was assessed using the Trait version of the Spielberger State/Trait Anxiety Inventory (STAI; Spielberger et al., 1983). This questionnaire is composed of 20 questions, assessing trait anxiety with questions such as “I lack self-confidence” and “I have disturbing thoughts”. Participants were asked to answer each statement using a response scale which runs from 1 = “Almost never” to 4 = “Almost always” in order to capture a stable dispositional tendency (trait) for anxiety.

#### 2.5.3. Beck Depression Index II (BDI)

Symptoms and severity of depression were evaluated using the BDI (Beck, Steer, Ball, & Ranieri, 1996). Participants responded to 21 questions designed to assess the individual’s level of depression (e.g. Sadness, pessimism, past failure etc.). The BDI items are scored on a scale from 0–3. All items were then summed for a BDI total score.

#### 2.5.4. Alcohol Use Questionnaire (AUQ)

The AUQ (Mehrabian & Russell, 1978) is a 15-item scale measuring in a detailed way the quantity of alcohol consumption (alcohol units of 8g drunk per week). For the past six months, participants were asked to estimate the number of drinking days, the usual quantity consumed and drinking pattern. For the purpose of our study, we used only the drinking quantity (i.e. alcohol units per week).

### 2.6. Procedure

The study was conducted at the Clinical Imaging Science Centre in Brighton, United Kingdom. Participants were asked to abstain drinking 24h before the experiment and gave fully informed consent. Demographic data (e.g. age, education level) were recorded and psychometric questionnaires were administered. Participants were breathalyzed and a urinary sample was collected to test for drug use. Task instructions were explained again and an inflatable Finometer finger cuff was then fitted to participants’ middle finger. After a 5 min recording calibration period, the participant was invited to start the Sternberg Task (Gendolla & Silvestrini, 2011). Each trial began with a 1000 ms fixation cross, followed by the EFE (20 ms) and a backward Mondrian mask (125 ms). This rapid series of events was immediately followed by a string of seven letters that remained on screen for 750ms, followed by a backward mask of seven “X” letters (750 ms). Then a target letter appeared at the centre of the screen until participants made a decision (max 2000ms), denoting whether or not the target letter was present in the earlier letter-string by pressing the right or the left arrow key, respectively. Next, a visual analogue scale allowed participants to rate their confidences for each trial, from “zero” to “extreme” (3000ms). In case of non-response, the message “Please answer more quickly” was presented during 3000ms. Finally, an inter stimuli interval of randomised duration (between 100 to 300 ms) was added before the beginning of the next trial. The maximum duration of a trial was 7s 645ms. Participants were encouraged to answer as quickly and accurately as possible. Reaction times (RT) and accuracy were recorded. The order in which these blocks were presented was randomized. Pilot study data showed participants would be unaware of the emotional nature of the prime stimulus (Gendolla & Silvestrini, 2011). However this was not formally confirmed for each individual trial in the main study.

### 2.7. Data analyses

#### 2.7.1. Behavioural data processing

One participant was excluded due to a depression score above three standard deviations of the mean. Reaction time on the Sternberg task and accuracy of response were recorded for each trial. Very fast reaction times (< or = 100ms) were deleted (Whelan, 2008). Missing data were quantified. Seventy-six reaction times data points were missing on a total of 1240 observations. Rather than deleting these cases, we performed multiple imputation using predictive mean matching. Predictive mean matching (PMM) is a semi-parametric imputation approach which imputes missing values by means of the nearest-neighbour donor with a distance based on the expected values of the missing variables conditional on the observed covariates (Little, 1988) We calculated these using the Multiple Imputation by Chained Equations (MICE) package in R, with PMM as method of imputation and the number of imputations set at 5 (Van Buuren & Groothuis-Oudshoorn, 2011). Imputed data were checked and included in the dataset.

#### 2.7.2. Physiological data processing

Inter-beat-Interval (IBI, ms), beat-to-beat values of systolic blood pressure (mmHg; from Finapres) and event-related information were extracted from recordings in Spike. Physiological data were smoothed using a Gaussian function (set to 1) to create a constant signal over systolic peaks and average across potential spike artefacts. Events data were aligned and binned at 100□Hz. All data were exported to Matlab (MATLAB and Statistics Toolbox Release 2016a, The MathWorks, Inc., Natick, Massachusetts, United States). Trial-by-trial systolic blood pressure levels values were derived from the averaged height of systolic peaks between the EFE presentation and the end of the VAS presentation over each trial.

### 2.8. Statistical Analyses

#### 2.8.1. Correlations

Mean and standards deviations were computed for reaction time, accuracy and systolic blood pressure. Physiological and behavioral measures were correlated to psychometric data, using 2-tailed nonparametric correlations.

#### 2.8.2. Mixed effects linear models

We used mixed-effects modeling to test effects of the variables (accuracy, reaction times (RT) and systolic blood pressure (SBP)), measured on trial by trial basis, (Barr, Levy, Scheepers, & Tily, 2013).

Accuracy was analyzed using a generalized linear mixed model as the outcome was binary (binomial family; Inaccurate =0; Accurate =1). To satisfy normality assumption, reaction times were also analyzed using a generalized linear mixed-effects model (Lo & Andrews, 2015). After fitting different density to the observed reaction times distribution, the relative quality of the models was estimated using Akaike information criterion (AIC). The lower AIC (e.g. best fit) was observed when a Gamma distribution was fitted to the observed reaction time distribution.

The same basic model was tested for each of the two outcomes (i.e. accuracy and reaction time). The basic model included systolic blood pressure, emotion (3 levels: Neutrality=0; Sadness=1; Anger=2), TAS score and the interactions terms as predictors. Therefore, intercept reflects the outcome value in the neutral condition. Given the established influence of age on blood pressure reactivity, age was included in the basic model as a control variable. Finally, participants were specified as a random (subject) factor, allowing for random intercepts.

The basic model was then compared to a similar model that also included anxiety, depression and alcohol intake to test confounding effects of these variables, contrasting the goodness of fit of the models using likelihood ratio tests.

All continuous predictors were mean-centered prior being entered in models. Analyses were undertaken using the lme4 package (Bates, Maechler, Bolker, & Walker, 2015). For models including a random term, the default lme4 optimizer was used. Finally, *p* values were computed using lmerTest package (Kuznetsova, Brockhoff, & Christensen, 2014). All analyses were run in the R environment (version 3.4.2; RCoreTeam, 2013).

#### 2.8.3. Post-hoc analyses: Heart rate variability

In order to explore if accuracy, reaction times and systolic blood pressure were related to a deceleration at the heart rate level, we analysed the heart rate variability in the frequency domain. To do so, we used the software HRVAS (Ramshur,, http://sourceforge.net/projects/hrvas/?source=navbar). The Lomb-Scargle method was preferred as this method provides power spectral density estimates of unevenly sampled data (Laguna, Moody, & Mark, 1998). For each participant, we computed mean interbeat interval, low cardiac frequencies percentage (0.04Hz to 0.15Hz), high cardiac frequencies percentage (0.15Hz to 0.4Hz) and the ratio low-to-high cardiac frequencies. Finally, mean and standards deviations were computed for each variable. Physiological measures were correlated to behavioural data, using 2-tailed nonparametric correlations.

## 3. Results

### 3.1. Mean, standard deviations, correlation and sample characterisation

Means, standard deviations, ranges, and correlation coefficients between psychometric, physiological and behavioral measures, are presented in **Error! Reference source not found**.. We did not observe suprathreshold correlations across psychometric data, behavioral and physiological measures. Moreover, in this sample, we found also no positive correlation between alexithymia and resting systolic blood pressure. Concerning alexithymia scores, 10 participants were characterized as non alexithymic (32.26%), 10 participants were classified as intermediate (32.26%), 11 participants were classified as alexithymic (35.48%).

### 3.2. Accuracy

Accuracy was analyzed with emotion (3 levels: Neutrality=0; Sadness=1; Anger=2), systolic blood pressure, TAS and their interactions as fixed factors. The participant (subject) variable was defined as a random factor (see methods section). The model controlled for age. The distribution was set as binomial (see **Error! Reference source not found**.).

There was a main effect of emotion; anger primes elicited increased accuracy compared to sadness and neutrality conditions (β = -0.50, SE = 0.13, *p* < .001; see Figure 1). A main effect of systolic blood pressure was also observed: increased systolic blood pressure was further also related to increased accuracy (β = 1.56, SE = 0.6, *p* < .050). Interaction between systolic blood pressure and emotion (anger vs neutrality) was observed as a trend: Here, low systolic blood pressure was associated with better accuracy in the anger condition compared to the neutral condition. However, increased blood pressure led to similar accuracy in both conditions (β = -1.14, SE = 0.64, *p* = .070; see Figure 2). Interaction between TAS-20 and systolic blood pressure was also observed as a trend: While in non-alexithymic individuals, performance accuracy correlated positively with systolic blood pressure, alexithymic individuals showed increased accuracy at lower systolic blood pressures and their gain in accuracy with increasing systolic blood pressure was reduced compared to non-alexithymic individuals (β = -0.1, SE = 0.05, *p* =.058; see Figure 3). There was furthermore a main effect of age (β = -0.06, SE = 0.03, *p* < .050). Addition of control variables (anxiety, depression, alcohol intake) to the model did not significantly improve the goodness of fit (Basic model: AIC =2586.9; Model with covariates: AIC = 2590.4; comparison: χ^2^(3) = 2.53; *p* = .470).

**Figure 1:**
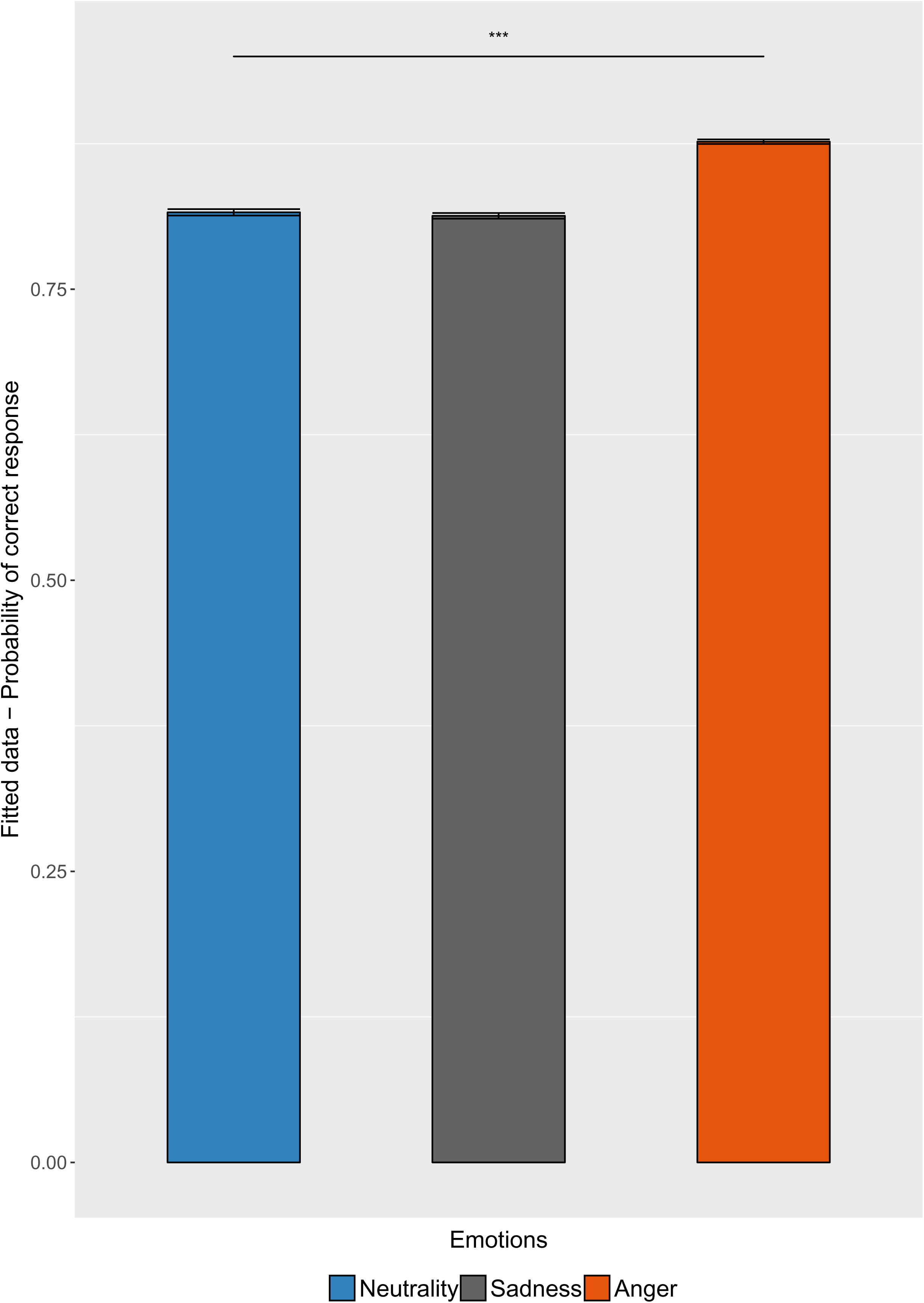
Main effect of emotion on probability of being accurate (*** *p* < .001)

**Figure 2:**
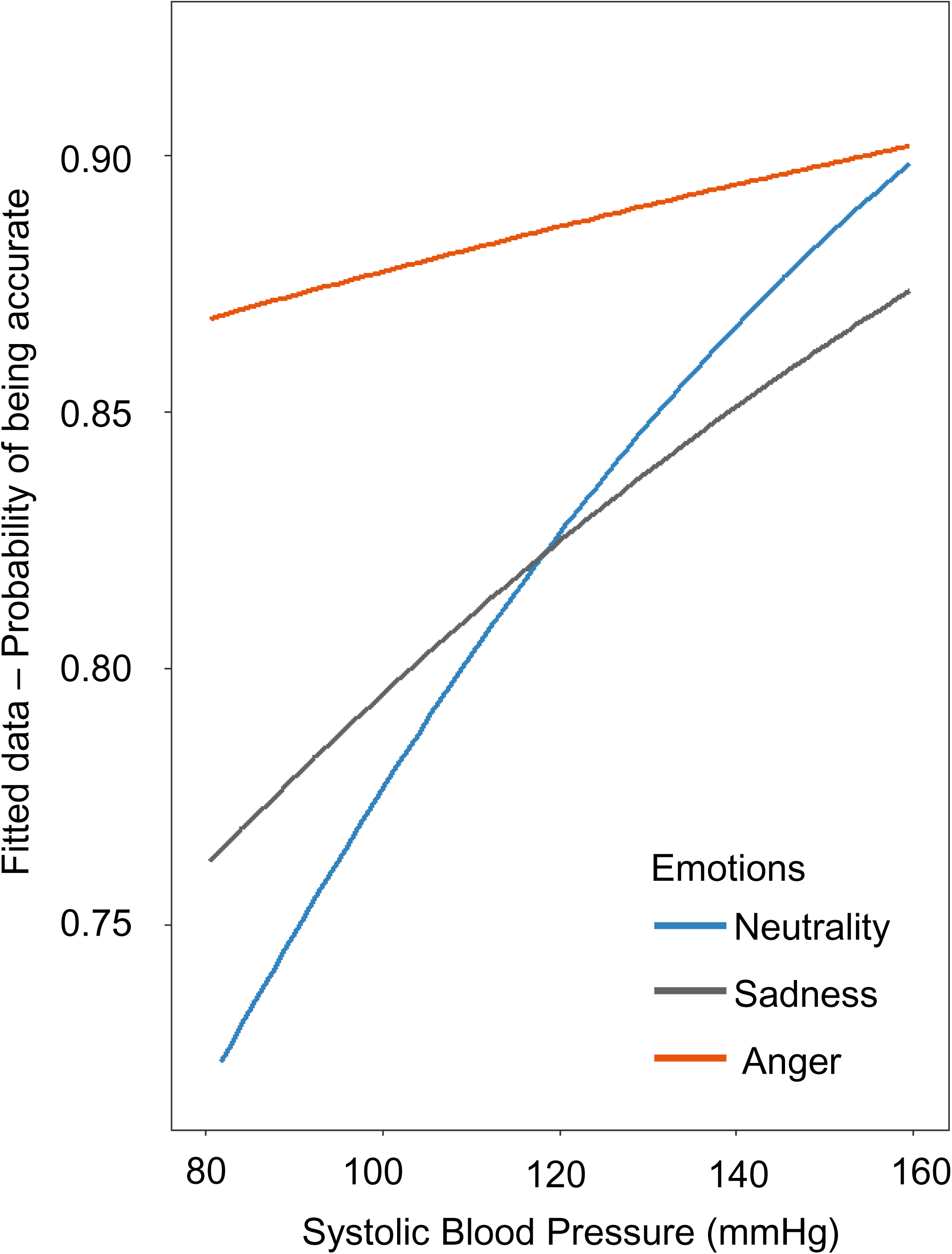
Trend of interaction between systolic blood pressure and emotion on probability of being accurate (*p* = .070)

**Figure 3:**
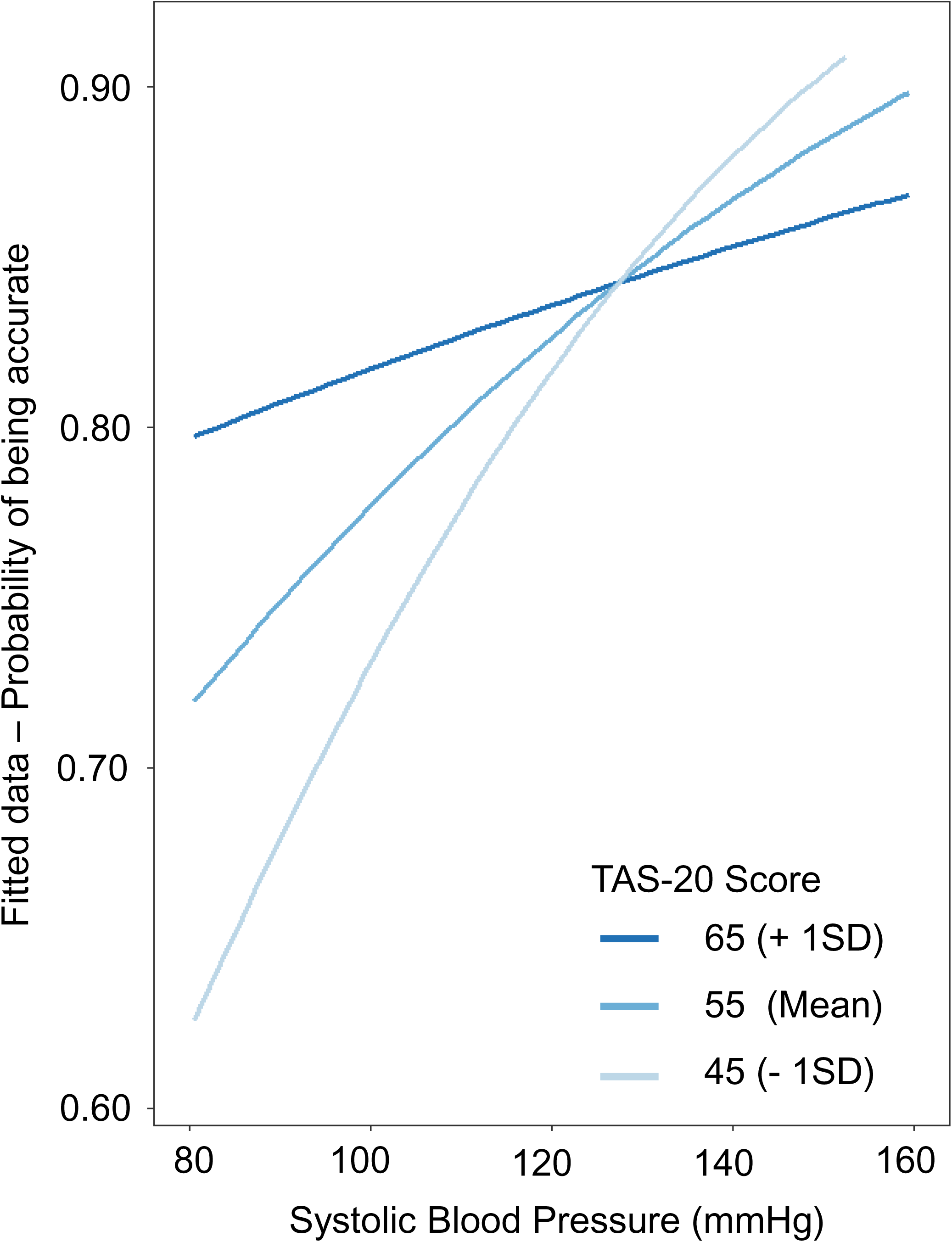
Trend of interaction between systolic blood pressure and Alexithymia (TAS-20 scores) on probability of being accurate (*p* = .058). To illustrate this interaction, high alexithymic (represented by Mean TAS-20 score + 1SD = 65), intermediate (represented by Mean TAS-20 score = 55) and non alexithymic (represented by Mean TAS-20 score -1SD = 45) were plotted.

### 3.3. Reaction Time

Reaction times were analyzed with emotion (3 levels: Neutrality=0; Sadness=1; Anger=2), systolic blood pressure, TAS and their interactions as fixed factors. The participant variable was defined as a random factor (see methods section above). The model controlled for age. The distribution was set as gamma. (see **Error! Reference source not found**.).

There was an interaction between systolic blood pressure and emotion (sadness vs. neutrality conditions): For neutral (and anger) primes, increases in systolic blood pressure was associated with prolongation of reaction times. However, increases in blood pressure during the sadness prime condition were associated with faster reaction times (β = -0.20, SE = 0.06, *p* < .001; see Figure 4). A main effect of systolic blood pressure on reaction time was observed as a trend (β = -0.14, SE = 0.08, *p* = .090). Addition of control variables (anxiety, depression, alcohol intake) to the model did not significantly improve the goodness of fit (Basic model: AIC =736.17; Model with covariates: AIC = 741.50; comparison: χ^2^(3) = 0.67; *p* = .880).

**Figure 4:**
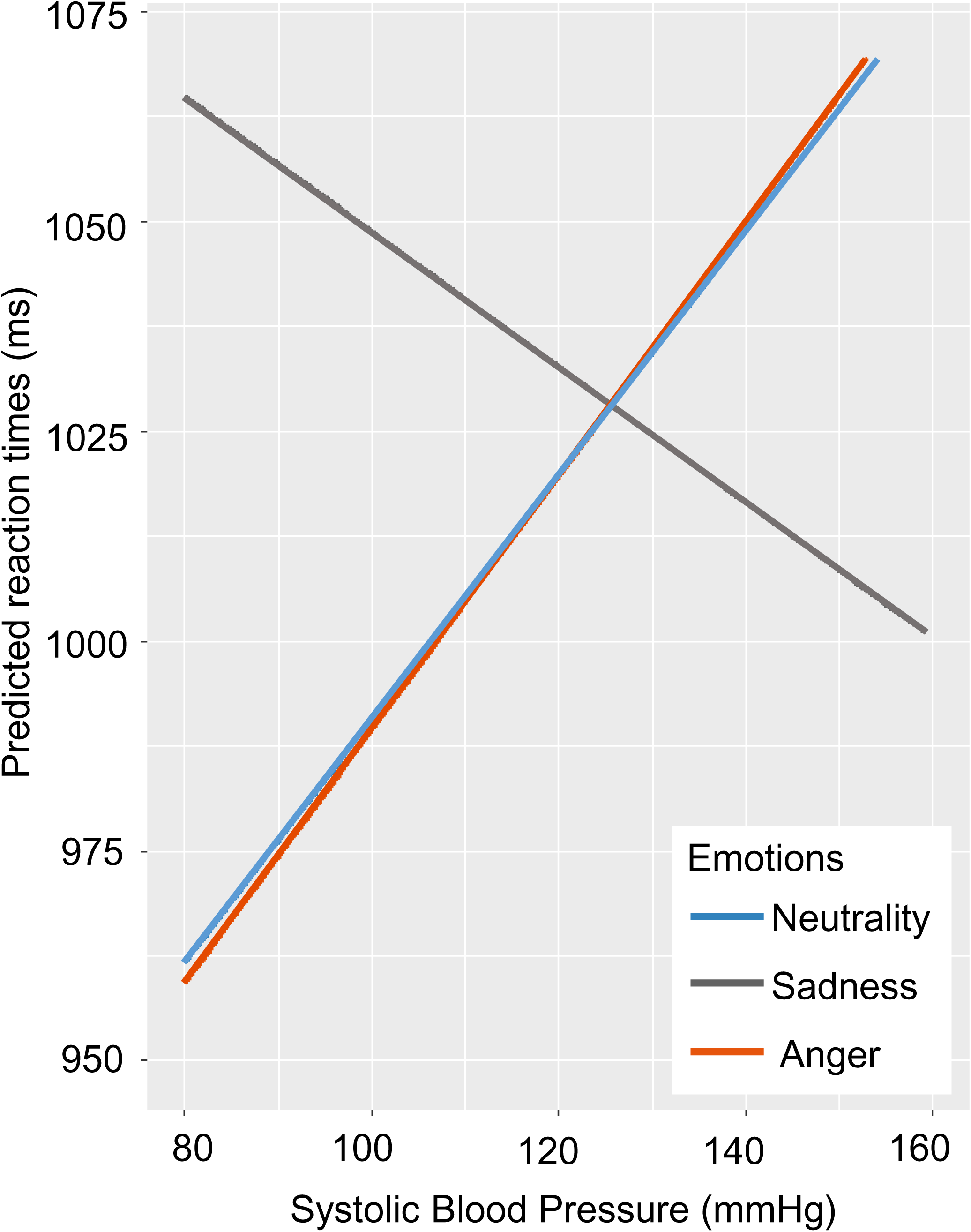
Interaction between systolic blood pressure and emotion on reaction times (*p* < .001)

### 3.4. Post-hoc analyses

Means, standard deviations, and ranges, as well as uncorrected correlation coefficients between physiological and behavioral measures, are presented in Table 4. We observed a negative correlation between mean interbeat interval and correct responses percentage: greater accuracy was associated with increased heart rate (τ= -.323, *p* = .012). Reaction times were negatively correlated with low frequencies percentage suggesting an association between shorter reaction times and increased sympathetic (or mixed) activity (τ= -.254; *p* = .045). We did not observe suprathreshold correlations across HRV-related measures and systolic blood pressure.

**Table 1:**
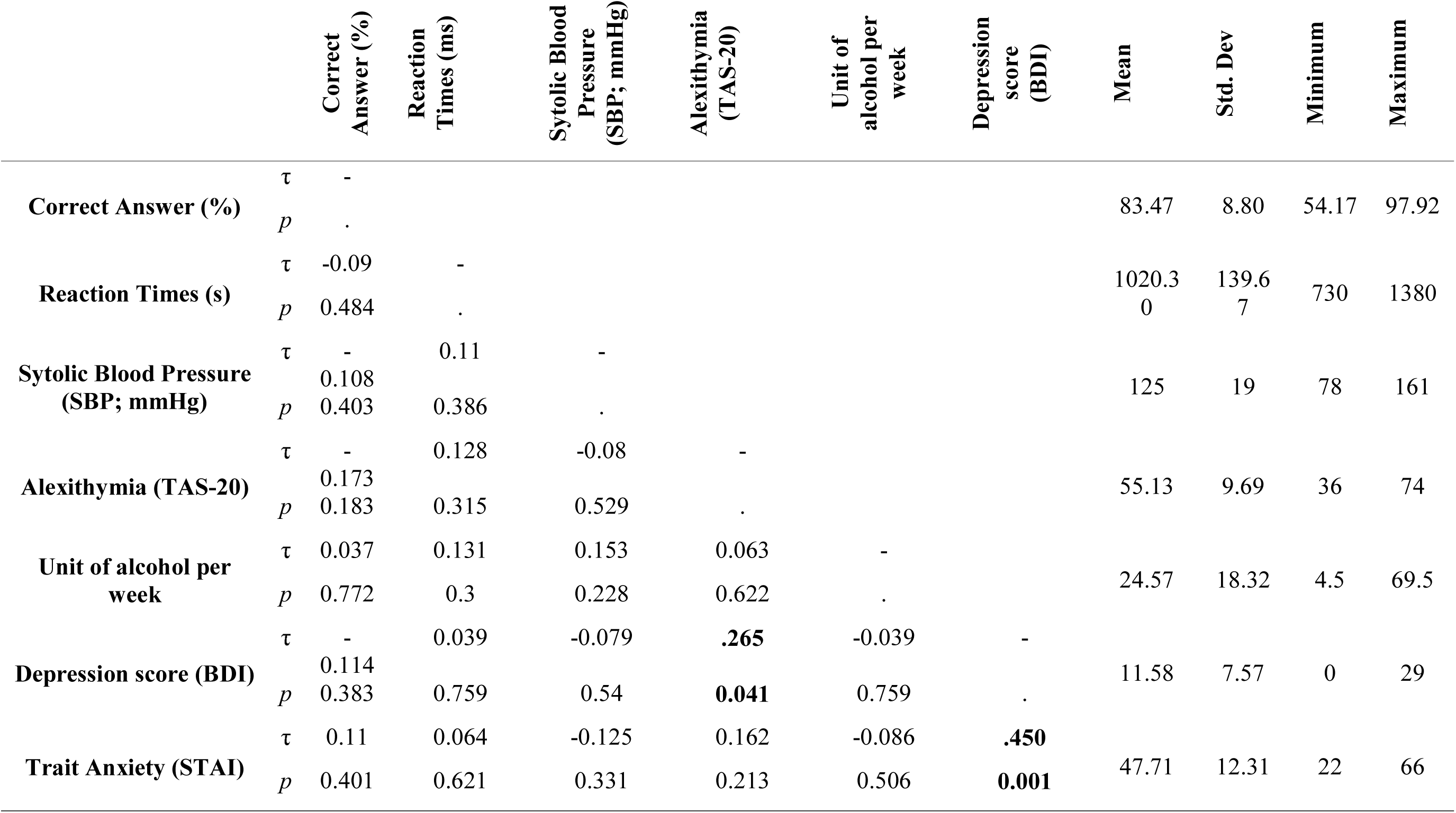
Mean, standard deviations, range as well as uncorrected Kendall’s tau correlation coefficients for psychometric, physiological and behavioural measures

**Table 2:**
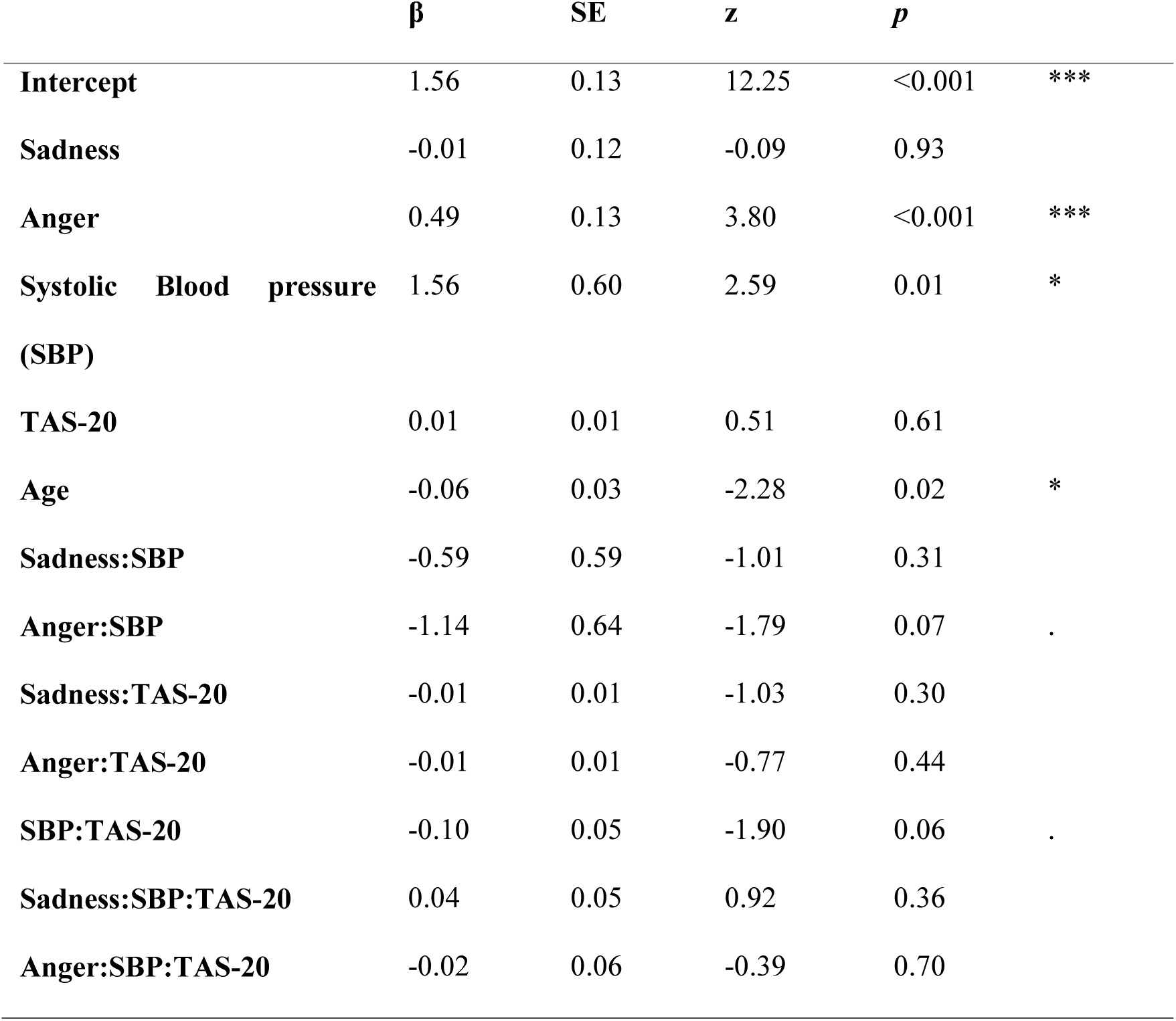
Mixed-effects regression model to explain accuracy, using Emotion (Neutrality, Sadness, Anger), systolic blood pressure (SBP) alexithymia (TAS-20) and their interactions as predictors, and, age as control variable. signif. codes: 0 ‘***’ 0.001 ‘**’ 0.01 ‘*’ 0.05 ‘.’ 0.1

| | $\beta$ | SE | $z$ | $p$ | |
| --- | --- | --- | --- | --- | --- |
| **Intercept** | 1.56 | 0.13 | 12.25 | <0.001 | *** |
| **Sadness** | -0.01 | 0.12 | -0.09 | 0.93 |  |
| **Anger** | 0.49 | 0.13 | 3.80 | <0.001 | *** |
| **Systolic Blood pressure (SBP)** | 1.56 | 0.60 | 2.59 | 0.01 | * |
| **TAS-20** | 0.01 | 0.01 | 0.51 | 0.61 |  |
| **Age** | -0.06 | 0.03 | -2.28 | 0.02 | * |
| **Sadness:SBP** | -0.59 | 0.59 | -1.01 | 0.31 |  |
| **Anger:SBP** | -1.14 | 0.64 | -1.79 | 0.07 | . |
| **Sadness:TAS-20** | -0.01 | 0.01 | -1.03 | 0.30 |  |
| **Anger:TAS-20** | -0.01 | 0.01 | -0.77 | 0.44 |  |
| **SBP:TAS-20** | -0.10 | 0.05 | -1.90 | 0.06 | . |
| **Sadness:SBP:TAS-20** | 0.04 | 0.05 | 0.92 | 0.36 |  |
| **Anger:SBP:TAS-20** | -0.02 | 0.06 | -0.39 | 0.70 |  |

**Table 3:** Mixed-effects regression model to explain reaction times, using Emotion (Neutrality, Sadness, Anger), systolic blood pressure (SBP) alexithymia (TAS-20) and their interactions as predictors, and, age as control variable. signif. codes: 0 ‘***’ 0.001 ‘**’ 0.01 ‘*’ 0.05 ‘.’ 0.1

| | $\beta$ | SE | $z$ | $p$ | |
| --- | --- | --- | --- | --- | --- |
| **Intercept** | 1.01 | 0.03 | 30.08 | <0.001 | *** |
| **Sadness** | -0.01 | 0.01 | -1.03 | 0.30 |  |
| **Anger** | 0.00 | 0.01 | -0.07 | 0.95 |  |
| **Systolic Blood pressure (SBP)** | -0.14 | 0.08 | -1.69 | 0.09 | . |
| **TAS-20** | 0.00 | 0.00 | -0.51 | 0.61 |  |
| **Age** | 0.00 | 0.01 | 0.26 | 0.79 |  |
| **Sadness:SBP** | 0.20 | 0.06 | 3.36 | <0.001 | *** |
| **Anger:SBP** | -0.01 | 0.06 | -0.19 | 0.85 |  |
| **Sadness:TAS-20** | 0.00 | 0.00 | 0.94 | 0.35 |  |
| **Anger:TAS-20** | 0.00 | 0.00 | -1.01 | 0.31 |  |
| **SBP:TAS-20** | -0.01 | 0.01 | -1.23 | 0.22 |  |
| **Sadness:SBP:TAS-20** | -0.01 | 0.00 | -1.58 | 0.11 |  |
| **Anger:SBP:TAS-20** | 0.00 | 0.00 | 0.27 | 0.79 |  |

**Table 4:**
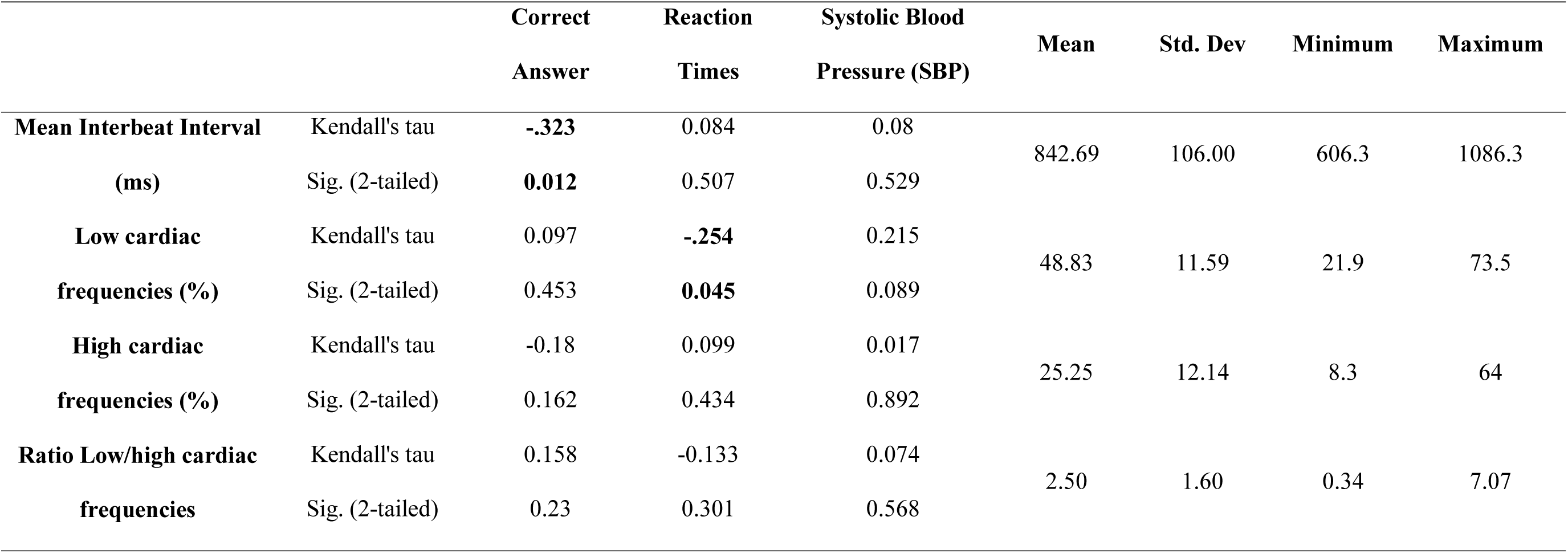
Mean, standard deviations, range as well as Kendall’s tau correlation coefficients for heart rate variability-related and behavioural measures.

|  |  | **Correct**<br>**Answer** | **Reaction**<br>**Times** | **Systolic Blood**<br>**Pressure (SBP)** | **Mean** | **Std. Dev** | **Minimum** | **Maximum** |
| --- | --- | --- | --- | --- | --- | --- | --- | --- |
| **Mean Interbeat Interval** | Kendall's tau | **-.323** | 0.084 | 0.08 |  |  |  |  |
| **(ms)** | Sig. (2-tailed) | **0.012** | 0.507 | 0.529 | 842.69 | 106.00 | 606.3 | 1086.3 |
| **Low cardiac** | Kendall's tau | 0.097 | **-.254** | 0.215 |  |  |  |  |
| **frequencies (%)** | Sig. (2-tailed) | 0.453 | **0.045** | 0.089 | 48.83 | 11.59 | 21.9 | 73.5 |
| **High cardiac** | Kendall's tau | -0.18 | 0.099 | 0.017 |  |  |  |  |
| **frequencies (%)** | Sig. (2-tailed) | 0.162 | 0.434 | 0.892 | 25.25 | 12.14 | 8.3 | 64 |
| **Ratio Low/high cardiac** | Kendall's tau | 0.158 | -0.133 | 0.074 |  |  |  |  |
| **frequencies** | Sig. (2-tailed) | 0.23 | 0.301 | 0.568 | 2.50 | 1.60 | 0.34 | 7.07 |

## 4. Discussion

In this present study, we tested how systolic blood pressure changes influence cognition during the processing of emotional primes, and explored how alexithymia might contribute to this relationship. We found an effect of emotional arousal on the accuracy of letter-string judgments: individuals were more accurate in their judgments when primed by angry faces compared to neutral faces. Moreover, better accuracy was associated with increased systolic blood pressure, regardless of the emotion. However, systolic blood pressure interacted also with the emotion quality of the prime on reaction time responses, where increasing blood pressure slowed reaction times following anger and neutrality primes but did not impact responses following sadness primes. We saw no direct moderating effect of alexithymia on accuracy or reaction times in our participants. Alexithymia did not correlate with blood pressure level. Nevertheless, a trend of interaction between systolic blood pressure and alexithymia on performance accuracy was still observed. Non-alexithymic individuals were more accurate in conditions associated with increased systolic blood pressure. In fact, alexithymic individuals were more accurate under low blood pressure compared to non-alexithymic individuals, whereas the inverse was observed under high blood pressure. Overall, alexithymic participants did not seem to benefit cognitively from blood pressure increases as much as non-alexithymic participants.

Our first main finding was an effect of arousal of the affective primes on decision-making accuracy. Here, participants were more accurate on the Sternberg task after being primed by angry faces compared to neutral faces. These results are coherent with existing literature: briefly flashed visual anger stimuli influence behavior, even without stimulating explicit affective responses (Gendolla, 2012; S. T. Murphy & Zajonc, 1993; Winkielman, Berridge, & Wilbarger, 2005). Anger is a negative-valenced and particularly salient emotion, which preferentially captures attentional resources (Burra, Barras, Coll, & Kerzel, 2016; Burra, Coll, Barras, & Kerzel, 2017; Feldmann-Wustefeld, Schmidt-Daffy, & Schubo, 2011; Hodsoll, Viding, & Lavie, 2011; Pinkham, Griffin, Baron, Sasson, & Gur, 2010; Shasteen, Sasson, & Pinkham, 2014). Presentation of angry face stimuli can increase visual short-term memory and working memory via modulation of basal ganglia activation (Jackson, Wolf, Johnston, Raymond, & Linden, 2008; Jackson, Wu, Linden, & Raymond, 2009). One potential explanation for increased accuracy after anger priming is the triggering of a hypervigilant state by the emotional anger prime, enhancing attentional deployment, which improves task performance. Alternatively, when primed by anger, the participants experienced a subjective reduction in task demand and a consequent increased ease in performance, when compared to the neutral and sadness priming conditions (Chatelain, Silvestrini, & Gendolla, 2016; Gendolla & Silvestrini, 2011). The latter is consistent with the coupling of anger to appetitive and approach motivational systems (Carver & Harmon-Jones, 2009; Russell, 2003). Angry facial expressions facilitate the generation of approach rather than avoidance motor responses (Wilkowski & Meier, 2010) and dynamic angry faces increase motor corticospinal excitability, mediating implicit and automatic responses to threat (Hortensius, de Gelder, & Schutter, 2016).However, here we did not find a main effect of emotion on reaction times.

Our second main finding, nevertheless, was a significant interaction between priming condition and systolic blood pressure. In both anger and neutral priming conditions increases in systolic blood pressure evoked increased reaction times. However, this linear relationship was absent in the sadness priming condition: reaction times did not seem to be modulated by systolic blood pressure changes. Moreover, in the sadness condition, lower systolic pressure was associated with longer reaction times. Blood pressure increases following verbal anger primes have been previously observed to predict a prolongation of reaction time on a lexical task (Garfinkel et al., 2016). The difference between these effects of anger and sadness primes parallels earlier findings: Sadness primes during an easy task can increase cardiovascular responses compared to anger primes, yet in a difficult task, the inverse pattern is found (Freydefont, Gendolla, & Silvestrini, 2012). Moreover, even masked sadness stimuli are associated with greater perceived difficulty and less ease when performing tasks (increased reactions times) and, therefore, increase the likelihood of disengagement when task demand becomes excessive (Silvestrini & Gendolla, 2011a, 2011c). However, our data did not show any reaction time prolongation during the sadness condition, as might be predicted by the implicit affect primes effort (IAPE) model of Gendolla as a sign of disengagement.

Our third main finding was that, on a trial-by-trial basis, task performance accuracy was related to increased systolic blood pressure. These data extend the existent literature. As shown by intra-arterial recordings, a sympathetic mechanism is implicated in engendering the blood pressure increases that accompany simple reaction-time tasks, (Obrist et al., 1974; Paller & Shapiro, 1983). Lower blood pressure correlates with poorer performance on a visuospatial attentional task in young hypotensive women (Cellini, Covassin, de Zambotti, Sarlo, & Stegagno, 2013; Wharton et al., 2006). Moreover, pharmacological elevation of blood pressure improves cognitive performance in hypotensive patients (Duschek, Hadjamu, & Schandry, 2007). Typically a rise in blood pressure activates arterial baroreceptors, which ultimately inhibit both cardiac and cortical activity, impacting cognitive processes (Kimmerly, 2017; Rau, Pauli, Brody, Elbert, & Birbaumer, 1993). Natural or artificial baroreceptor stimulation can inhibit somatosensory afferent information flow (including pain) (Angrilli, Mini, Mucha, & Rau, 1997; Gray et al., 2010). This cardiac afferent mechanism is postulated to reduce input from the external environment and, thereby reduce inattention and distractibility. Correspondingly, our data (i.e. higher systolic blood pressure, higher accuracy), accompanied possibly by heartrate deceleration (reducing afferent cardiac feedback to the brain and thereby limiting interference with cognitive processes) might have been broadly in line with the notion of an increase in attentive observation of the environment (Lacey & Lacey, 1970). Speculatively, our observed increased blood pressure could have regulated cardiac output - via vagal influences. This control might have thus facilitated sensorimotor performance (Cellini et al., 2013; G. Park & Thayer, 2014; G. Park, Vasey, Van Bavel, & Thayer, 2013). Indeed, cardiac deceleration is itself modulated by arousal (e.g. threat) supporting the hypothesis of an evolutionary survival strategy (Hare, Wood, Britain, & Shadman, 1970; Libby, Lacey, & Lacey, 1973). However, we are cautious as we only observed a statistical trend in the interaction between blood pressure and emotion on performance accuracy. For the neutral condition, increased systolic blood pressure was associated with greater accuracy. First, we recognised the relevance of this observation to Lacey and Lacey’s (1970) hypothesis. This model describes an association between cardiac deceleration and attention (or ‘intake’) directed to stimulations from the external environment; congruent with the lower-level inhibitory influence on sensory processing and cortical excitability induced by baroreceptor activation. To test the Lacey and Lacey’s hypothesis, we conducted post-hoc analyses. These showed that greater accuracy and shorter reaction times were associated with faster heart rate and an increase in power of low frequency heart rate variability, respectively. Given the absence of an association between heartbeat deceleration or increased parasympathetic activity index (e.g. high cardiac frequencies) and accuracy, our data do not seem to support Lacey and Lacey’s hypothesis. Instead, the rise in systolic blood pressure and the greater accuracy seem to be both driven by increased arousal induced by affective priming. This interpretation is congruent with the observed increased accuracy, in the absence of a modulation bodily state, under anger priming condition. Moreover, this discrepancy between different emotional conditions suggests the involvement of emotion-specific pathways (Brooks et al., 2012; Lacey & Lacey, 1970). Also, it is important to specify that regression analyses do not allow determining whether the association between performance and physiological reactivity reflects a causal influence of the later to the former. For example, task performance and physiological activity might be both independent but parallel consequences of effortful cognitive processes, as proposed by the IAPE model (Gendolla, 2012).

A secondary aim of the study was to characterise the impact of alexithymia on the relationship between bodily changes and behaviors. Here, we found a trend of an interaction between alexithymia and systolic blood pressure. Compared to non-alexithymic, alexithymic individuals benefitted least from systolic blood pressure fluctuations. Alexithymia is classically characterized by increased blood pressure and reduced interoceptive abilities (Betka et al., 2017; Bornemann & Singer, 2017; Brewer et al., 2016; Gage & Egan, 1984; Jula et al., 1999; J. Murphy et al., 2017; Todarello et al., 1995). Given their atypical autonomic profiles, one could postulate that alexithymic people might also attribute less salience to bodily changes and show impaired integration of autonomic information when compared to non-alexithymic. Compensatory strategies developed by alexithymic individuals may explain, in part, why they are not impaired on the task. For example, alexithymic individual might use information related to bodily actions (e.g. increased somatosensory and motor areas activation) rather than affective states to correctly label emotional faces (Ihme et al., 2014). Alexithymic individuals may have particular difficulty in processing and automatically using high arousal emotional information in the context of cognitive challenges (Vermeulen et al., 2006). In that way, our data also suggest reduced integration of highly relevant emotional information in alexithymia. Further studies should clarify this relationship.

### 4.1. Limitations

Finally, we recognize limitations of our study. A larger sample might have increased statistical power and sensitivity to explore the impact of inter-individual characteristics on these relationships between body and behaviour. For example, it would have been interesting to measure trait anger and hostility, which is a factor known to modulate the effects of subliminal anger primes (Garfinkel et al., 2016; Wilkowski & Robinson, 2008). Another limitation is that after calibrating the stimuli independently, we did not conduct an awareness check during the task to establish the degree to which the very brief (20ms) primes were processed unconsciously by each partcipant. We, therefore, cannot guarantee that our affective prime stimuli were rendered fully subliminal (van der Ploeg et al., 2017), although this would be unusual for our rapid presentation of the primes. Also, an objective measure of effort mobilization would have permitted us to interpret our results more confidently in the framework proposed by the IAPE model (Gendolla, 2012). Lastly, recent studies highlight the importance of taking in account both affective and cognitive dimensions of alexithymia as their autonomic signatures may differ (Cecchetto, Rumiati, & Aiello, 2017; Martínez-Velázquez, Honoré, de Zorzi, Ramos-Loyo, & Sequeira, 2017). We did not have this degree of granularity within the present dataset.

### 4.2. Conclusion

In conclusion, our data demonstrate the interacting effects of peripheral autonomic changes and affective states in guiding mental processes. A growing literature highlights atypical autonomic profiles across a wide range of psychiatric disorders. Future studies should, therefore, clarify the weight given to autonomic information in the generation of behavioural responses, in vulnerable populations.

## Supporting information

Suuplementary Materiel

## Abbreviations

AIC: Akaike Information Criterion
AUQ: Alcohol Use Questionnaire
EFE: Emotional Facial Expression
IAPE: Implicit Affect Primes Effort (IAPE)
PMM: Predictive Mean Matching
SBP: Systolic Blood Pressure
TAS-20: Toronto Alexithymia Scale-20 items.

## Authors contribution

SB, SG, DD, HS and HDC were responsible for the study concept and design. SB and GP contributed to data acquisition. SB, DW and HDC assisted with data analysis and interpretation of findings. SB drafted the manuscript. DD, HS and HDC provided critical revision of the manuscript for important intellectual content. All authors critically reviewed content and approved final version for publication.

