## Supplementary material for "Relationship between systolic blood pressure and decision-making during emotional processing": Suuplementary Materiel: Supplementary Material.docx

List of stimuli from the Karolinska Directed Emotional Faces resource (KDEF <http://www.kdef.se/> Lundqvist, D., Flykt, A., & Öhman, A. (1998). The Karolinska Directed Emotional Faces - KDEF, CD ROM from Department of Clinical Neuroscience, Psychology section, Karolinska Institutet, ISBN 91-630-7164-9.

AF01ANS

AF01NES

AF01SAS

AF02ANS

AF02NES

AF02SAS

AF03ANS

AF03NES

AF03SAS

AF04ANS

AF04NES

AF04SAS

AF05ANS

AF05NES

AF05SAS

AF06ANS

AF06NES

AF06SAS

AF07ANS

AF07NES

AF07SAS

AF08ANS

AF08NES

AF08SAS

AF09ANS

AF09NES

AF09SAS

AF10ANS

AF10NES

AF10SAS

AF11ANS

AF11NES

AF11SAS

AF12ANS

AF12NES

AF12SAS

AF13ANS

AF13NES

AF13SAS

AF14ANS

AF14NES

AF14SAS

AF15ANS

AF15NES

AF15SAS

AF16ANS

AF16NES

AF16SAS

AM01ANS

AM01NES

AM01SAS

AM02ANS

AM02NES

AM02SAS

AM03ANS

AM03NES

AM03SAS

AM04ANS

AM04NES

AM04SAS

AM05ANS

AM05NES

AM05SAS

AM06ANS

AM06NES

AM06SAS

AM07ANS

AM07NES

AM07SAS

AM08ANS

AM08NES

AM09ANS

AM09NES

AM09SAS

AM10ANS

AM10NES

AM10SAS

AM11ANS

AM11NES

AM11SAS

AM12ANS

AM12NES

AM12SAS

AM13ANS

AM13NES

AM13SAS

AM14ANS

AM14NES

AM14SAS

AM15ANS

AM15NES

AM15SAS

AM16ANS

AM16NES

AM16SAS

AM17SAS
